# Effects of Llama-Derived Hyperimmune Serum on Motility and Viability of *Echinococcus granulosus* Protoscoleces

**DOI:** 10.64898/2026.04.30.721736

**Authors:** María J. Fernández Salom, Mónica P.A. Carabajal, David Di Lullo, Héctor D. Villa Micó, Enrique Formentini, María del Rocío Cantero, Horacio F. Cantiello

**Affiliations:** Laboratorio de Canales Iónicos, Instituto Multidisciplinario de Salud, Tecnología y Desarrollo (IMSaTeD), CONICET-UNSE, Santiago del Estero, Argentina; Laboratorio de Farmacología y Toxicología, Facultad de Ciencias Veterinarias, Universidad Nacional del Litoral, Argentina

**Keywords:** Echinococcus granulosus, *Lama glama*, Protoscoleces, Camelid hyperimmune serum, Passive immunotherapy, Motility inhibition, Viability assays, ELISA, Cystic echinococcosis

## Abstract

Cystic echinococcosis (CE), caused by the larval stage of *Echinococcus granulosus*, remains a significant public health and veterinary problem in endemic regions. Although chemotherapy and control programs exist, the development of complementary immunotherapeutic tools is increasingly needed. This study evaluated the generation and functional activity of hyperimmune serum (HIS) produced in three adult male castrated llamas (*Lama glama*) immunized with antigenic material derived from protoscoleces (PSCs) of the parasite. Sera collected after each of the first six immunizations were assessed by ELISA to quantify antigen-specific IgG responses, and their biological effects were tested in vitro using viable PSCs. Motility was measured using video-assisted paired-image scoring across serial serum dilutions (1:2–1:2048), and the methylene blue exclusion assay was used to assess viability. Hyperimmune serum produced a clear, reproducible, dose-dependent inhibition of PSC motility and viability. Higher titers of early inoculations reduced motility by 70–85%, while sera from the fifth and sixth inoculations achieved complete suppression. Naïve serum and PBS controls showed no inhibitory effect. ELISA titers strongly correlated with biological activity, indicating that higher humoral responses predicted functional inhibition. These findings demonstrate the feasibility of generating potent anti-*Echinococcus granulosus* polyclonal antibodies in camelids and support their potential application in passive immunization strategies. The study establishes a foundation for future development of llama-derived immunobiological reagents, including nanobody-based tools, for the control of cystic echinococcosis.

## INTRODUCTION

Cystic echinococcosis, caused by the larval stage of *Echinococcus granulosus*, remains a major zoonotic disease in regions where livestock production is prominent, including large areas of South America (Romig 2021). Despite advances in diagnosis and treatment, reinfections and persistent transmission continue to generate substantial public health and economic burdens. (Larrieu 2019; Romig 2021) Chemotherapeutic interventions, although useful, have limitations, including incomplete efficacy, prolonged treatment courses, and variable parasite susceptibility. These challenges underscore the need for complementary or alternative strategies, including immunological approaches that disrupt parasite establishment, survival, or viability.

The protoscolex, the infective larval stage within hydatid cysts, is central to disease perpetuation. As a metabolically active organism lacking a digestive system, it relies on nutrient exchange across its syncytial outer covering, the tegument—a highly specialized interface that protects the parasite and mediates host–parasite interactions (Dalton 2004). However, immunological strategies directed at PSC surface antigens remain comparatively underexplored.

South American camelids such as llamas (*Lama glama*) provide a distinctive immunobiological platform for developing anti-parasitic tools. These species naturally produce conventional antibodies and heavy-chain–only antibodies (HCAbs), the latter of which give rise to single-domain fragments known as nanobodies (VHHs) (Hamers-Casterman 1993; Muyldermans 2013). Nanobodies exhibit high solubility, exceptional thermal and structural stability, and the ability to bind cryptic epitopes inaccessible to conventional antibodies (Muyldermans 2001, 2013). Since their discovery in camelid serum, nanobodies and their precursor polyclonal responses have emerged as promising reagents for targeting complex pathogens, including helminths. (Hussack & Tanha, 2010, Muyldermans, 2016).

Immunizing llamas with antigens derived from *E. granulosus*—including hydatid cyst fluid, somatic extracts, and PSC-derived proteins—offers an opportunity to generate hyperimmune sera enriched in antibodies capable of recognizing key parasite structures. Beyond serving as precursors for future nanobody development, these polyclonal antibodies may exert direct biological effects on PSC physiology.

The present study investigated whether hyperimmune serum produced in llamas can functionally impair PSC motility and viability in vitro. Using quantitative motility scoring, viability assays, and ELISA-based profiling of antigen-specific IgG responses, we examined the dose-dependent impact of these sera on parasite function. We explored correlations between antibody titers and biological inhibition. While electrophysiological studies were conducted in parallel as part of a broader research program, they fall outside the scope of this manuscript and will be reported separately. Here, we focus specifically on the relationship between immunization, humoral responses, and measurable functional effects on *E. granulosus* PSCs.

## MATERIALS AND METHODS

### Camelids Used for Experimentation

The study was conducted with llamas (*Lama glama*) reintroduced from the Jujuy highlands to the Ambargasta Salt Flats in the Santiago del Estero province, Argentina, in the town of Guanaco Sombriana (28.7584° S, 64.0715° O). This saline region, situated in the southwest of the province, covers approximately 4,200 km^2^ and is characterized by harsh environmental conditions for plant life, in which Chenopodiaceae species adapted to these ecosystems predominate (Figueroa 2009). The experimental herd consisted of three castrated adult male llamas, aged 2 to 3 years (Tampulli). Age was determined using the dental chronology of domestic South American camelids (Calle 1982, Bustinza 2001). The animals were treated with anthelmintic drugs based on the results of copro-parasitological analysis and received individual treatments for ectoparasites. The animals were also vaccinated against clostridial diseases.

The animals were fed on planted native pastures adapted to saline ecoregions (*Cenchrus spp*.) and native xerophytic species of the ecosystem, such as black sea lavender (*Allenrolfea vaginata*), white mesquite (*Prosopis alba*), black mesquite (*Prosopis nigra*), acacia (*Acacia aroma*), cardon cactus (*Echinopsis atacamensis*), and occasionally pastures of the genera Gramma rhodes (*Chloris gayana*), *Sporobolus*, and *Paspalum*. The native pastures are adapted to temperate and warm subtropical regions with seasonal rainfall and a prolonged dry season, characterized by their resistance to drought, intensive grazing, and burning. The average chemical composition of the native pastures was: dry matter 25.54 ± 3.11%, crude protein 10.42 ± 1.27%, neutral detergent fiber 65.2 ± 2.34%, and acid detergent fiber 35.65 ± 3.08%. Feeding was supplemented with a mixture of alfalfa (*Medicago sativa*) and crushed corn grain (*Zea mays L*.) at 1% of body weight during the winter, when natural pastures were scarce. Drinking water was administered *ad libitum*.

### Llama Immunization and Serum Collection

The animals were under continuous veterinary supervision and clinically evaluated before inclusion, ensuring the absence of clinical signs and seronegativity for *E. granulosus*, as confirmed by initial serological testing. Immunization was performed under field conditions, in compliance with national guidelines for the use of animals for experimental purposes. The general conditions of each animal were monitored throughout the protocol. The presence of local or systemic reactions to the inoculation was noted, and the entire clinical evolution was documented. No signs of pain, persistent inflammation, or significant behavioral changes were observed and recorded on the health observation forms.

For the vaccine formulation, the antigenic extract of *E. granulosus* (see below) was resuspended in sterile phosphate-buffered saline (PBS) and dosed at a concentration of 250 µg/mL. The vaccine emulsion was prepared in a 1:1 ratio with aluminum hydroxide as an adjuvant (1 mL of antigen per 1 mL of adjuvant) immediately before application, to ensure the integrity of the immunogenic proteins and the stability of the emulsion (Lindblad 2004).

The immunization protocol included a total of six subcutaneous inoculations (Fig. 1), applied in the lateral cervical region, distributed in two stages designed to maximize the production of polyclonal antibodies against a complex and unfractionated antigenic extract (Tizard 2018): 1) primary immunization phase, consisting of two initial doses separated by an interval of 15 days, intended to induce primary immunological sensitization, and 2) a booster phase, consisting of four additional doses administered with the same fortnightly periodicity, aimed at enhancing the humoral response through the booster effect.

**Fig. 1:**
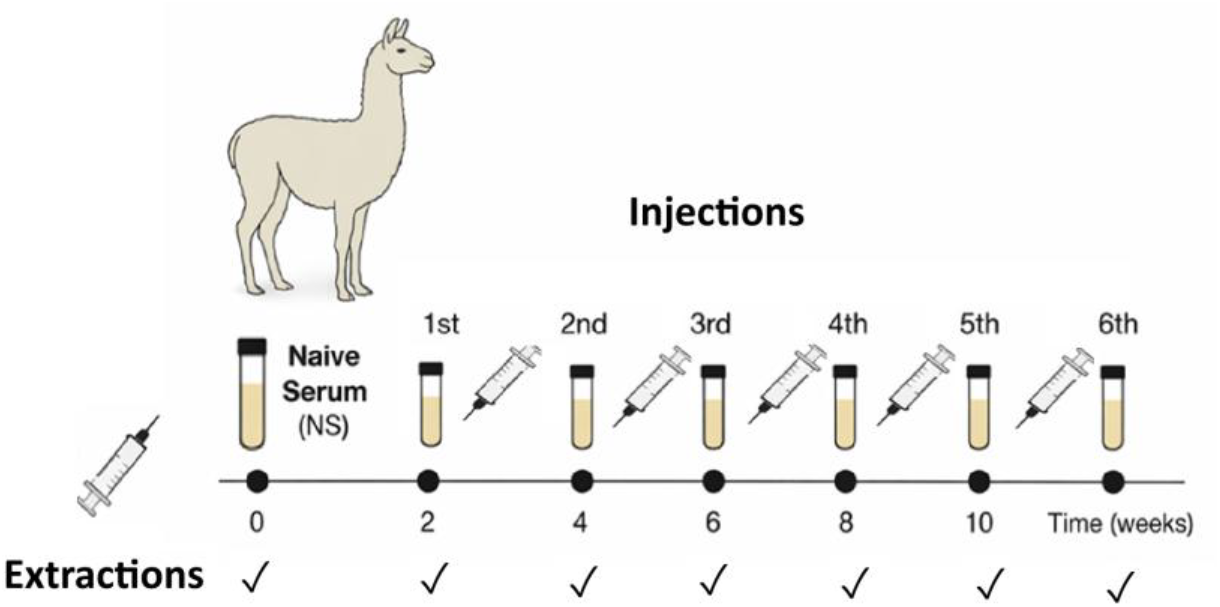
Immunization protocol and serum collection schedule. Llamas were subjected to six sequential subcutaneous immunizations with E. granulosus antigenic extract at 2-week intervals. The first two injections correspond to the primary immunization phase, followed by four booster doses. Blood samples were collected before each inoculation to obtain naïve and post-immunization sera for downstream analyses.

Before the first immunization, blood was drawn from each animal to obtain baseline (naïve) serum (NS), which was used as a negative control in the comparative trials. Subsequent blood draws were performed 24 h before each booster, using jugular venipuncture with a 20 mL syringe and an 18-gauge needle. The samples were collected in 15 mL Falcon tubes, allowed to clot at room temperature, and centrifuged at 3,000 rpm for 5 minutes. The serum was separated from the clot, aliquoted into 1.5 mL microtubes, and stored at –80°C until use.

The hyperimmune serum (HIS) from each llama was evaluated using an in-house-developed ELISA assay, which enabled selection of batches with the highest anti-*E. granulosus* antibodies, for later use in in vitro and in vivo assays.

### Source of Parasite Material

Protoscoleces (PSCs) of *E. granulosus* were obtained from hydatid cysts collected from naturally infected bovine livers and lungs at a local abattoir in Santiago del Estero, Argentina. Cysts were aseptically opened, and the parasitic material was filtered through sterile gauze to remove debris. The PSC suspension was washed three times with sterile PBS (pH 7.4) supplemented with antibiotics (penicillin 100 U/mL, streptomycin 100 µg/mL) and amphotericin B (0.25 µg/mL). The preparation was then gravity-sedimented for 30 min, and the supernatant was discarded to retain the viable PSCs.

Viability was assessed microscopically by methylene blue exclusion; only preparations with ≥90% viability were used for experiments. PSCs were maintained at 4°C and used within one month of collection. For functional assays, standardized suspensions containing approximately 3,000–5,000 PSCs/mL were prepared, as previously described (Carabajal 2023).

### Preparation of Antigenic Material from PSCs

Total antigenic extracts for use in camelid immunization were prepared from viable PSCs. The parasites were washed (3X) with sterile PBS (pH 7.4) to remove hydatid fluid residues and debris, then resuspended in lysis buffer at a ratio of 200 µL per ∼1,500 PSCs. Cell disruption was achieved by three freeze–thaw cycles (liquid nitrogen/37°C water bath), followed by mechanical sonication (three 30-s pulses at 40% power, with 60-second intervals on ice).

After lysis, samples were vortexed and centrifuged at 13,000*g* for 5 min at 4°C, and the lipid layer was discarded. Proteins were precipitated by adding cold acetone (−20°C) at a 2:1 acetone-to-extract ratio and incubating overnight at −20°C. The mixture was then centrifuged at 10,000*g* for 5 min, and the resulting pellet was air-dried and resuspended in sterile PBS. Extracts were aliquoted and stored at 4°C until use.

Protein concentration was determined using both Bradford and BCA assays, yielding typical ranges of 350–1000 µg/mL. For immunization, antigen solutions were adjusted to 250 µg/mL and emulsified 1:1 with aluminum hydroxide adjuvant (Imject™ Alum) immediately before administration to the camelids.

### Image Acquisition and Processing

Parasites were observed with a Zeiss® microscope using a 10× objective (NA 0.25), illuminated by a halogen lamp (12 V/100 W) with an intensity regulator; the diaphragm was adjusted for phase contrast when necessary. Images were captured with an Axocam (1920×1080 px) camera connected via USB 3.0 to a personal computer using the Micromanager 2.0 capture software and saved in uncompressed TIFF format. Micrographs were pseudo-colored solely for the presentation of results; all quantitative analyses were performed on 8-bit grayscale images. For each well, two sequential images were captured at 10-sec intervals under 50× magnification.

### Viability Assay

Viability was determined using the methylene blue exclusion method under bright-field illumination at 10× magnification. PSCs that absorbed the dye were classified as non-viable. In contrast, colorless PSCs showing spontaneous flame-cell activity and motility were considered viable (Fig. 2A). A viability threshold of ≥90% was required before initiating each experimental run. Viability scoring was performed in parallel with motility assays, but was analyzed independently to ensure robust quality control of the biological material used.

**Fig. 2:**
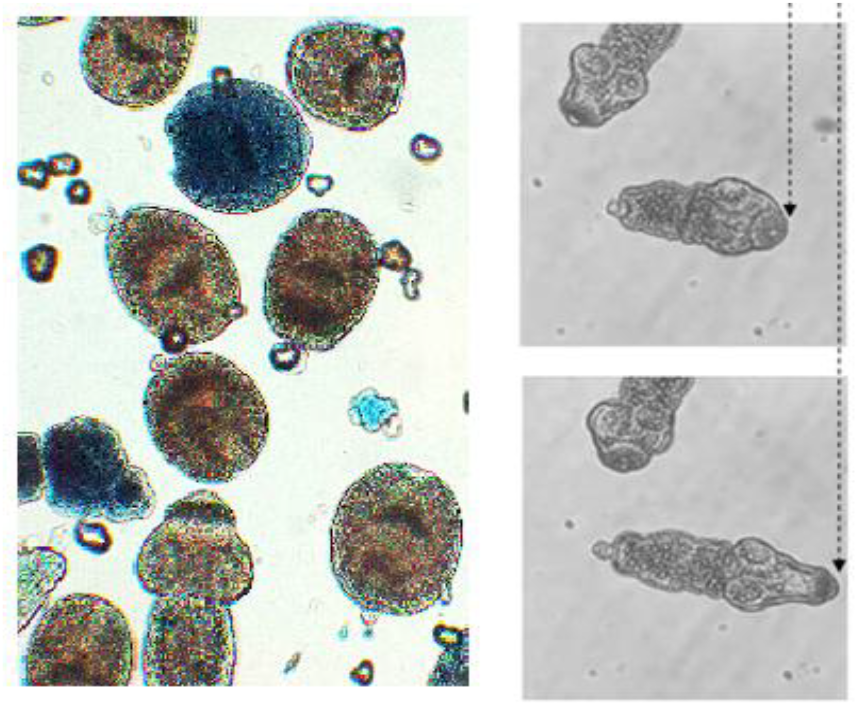
Assessment of PSC viability and motility. **Left**. Representative micrographs of PSCs stained with methylene blue. Viable PSCs exclude the dye, whereas non-viable PSCs exhibit dye uptake. **Right**. Quantification of PSC motility using paired-image analysis. Motility was defined as the percentage of PSCs exhibiting displacement between sequential images acquired under identical conditions.

### Motility Assay

The motility of *E. granulosus* PSCs was assessed using a quantitative, image-based, *in vitro* assay (Fig. 2B). PSCs were maintained in sterile Ringer’s Krebs solution (RKS) and pre-incubated for 24 h at 37°C in 5% CO_2_ to promote evagination and stabilize physiological activity. Assays were performed in 96-well flat-bottom microplates, each well receiving 10–30 PSCs in a final volume of 50 µL.

For each serum sample (HIS or naïve serum, NS), serial 1:2 dilutions were prepared across a horizontal row of wells (1:2 to 1:2024). Briefly, 25 µL of serum was added to the first well, and 25 µL PBS to wells 2–12. Serial dilutions were generated by transferring 25 µL sequentially across the row, discarding the final 25 µL from well 11, leaving well 12 as the PBS-only control. PSCs were then added (25 µL/well), yielding a final serum proportion of 50% per dilution step. Microplates were covered and incubated at 37°C for 24 and 48 h. Parasite motility was estimated by analyzing TIF images using Windows Image Viewer. The movement of evaginated and invaginated parasites, or motility data, was determined by comparing two images taken 10 seconds apart for each sample. The total number of parasites observed, the number exhibiting movement, and the number of evaginated and invaginated parasites were recorded. (Fig. 2). Motility was expressed as the percentage of motile PSCs with respect to the total number of parasites within the analyzed region. Motility was shown as the mean ± SEM percentage of mobile PSCs per field.

### ELISA Assay

Antibody responses in immunized llamas were quantified using an indirect ELISA. High-binding 96-well plates were coated overnight at 4°C with *E. granulosus* antigen (1–5 µg/mL) in carbonate or PBS buffer (pH 7.4–9.6). After antigen adsorption, wells were washed (x3) with PBST (PBS + 0.05% Tween-20) and blocked for 30 min at room temperature with 5% BSA in PBST. Serum samples were added in serial 1:2 dilutions (typically 1:2 to 1:512) prepared in PBST containing 1% BSA, and plates were incubated for 60 min at room temperature with gentle agitation. After washing, HRP-conjugated goat anti-llama IgG (H+L) was added at a 1:1000–1:10,000 dilution and incubated for 60 min. Following the final washes, TMB substrate (Thermo cat#34,029) was added, and the reaction was stopped with 0.16 M H_2_SO_4_. Absorbance was recorded at 450 nm using a Thermo Multiskan FC reader.

### Statistical Analysis and Data Fitting

Data were expressed as Mean ± SE. Comparisons between groups were performed using one-way ANOVA followed by Tukey’s post-hoc test. Correlations between ELISA titers and biological activity (motility, viability) were assessed using Pearson’s correlation coefficient. A *p*-value < 0.05 was considered statistically significant. Analyses were conducted using GraphPad Prism 9.

To explore serum potency, motility data were fitted using two complementary models. First, a standard Hill-type model, as follows:

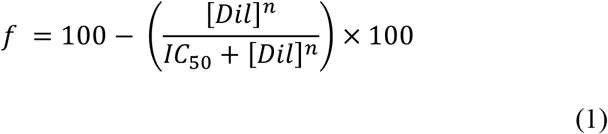

where [*Dil*] is the serum dilution used, *IC*_50_ is the half-maximal inhibitory concentration, and *n* is the Hill coefficient.

In addition, to account for potential heterogeneity in antibody–target interactions, the data were also fitted to a two-site model assuming independent binding sites with different affinities:

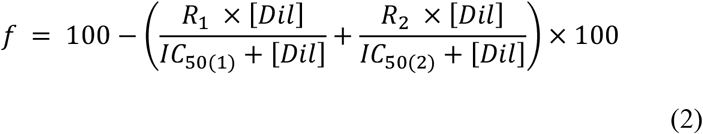

Where *R*_1_ and *R*_2_ are the maximal serum activities for each component [*Dil*] is the serum dilution used, and *IC*_50(1)_ and *IC*_50(2)_ are the half-maximal inhibitory concentrations for the low and high affinity sites.

ELISA data were fitted with an adaptation of equations (1) and (2), such that

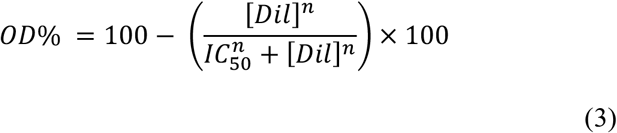

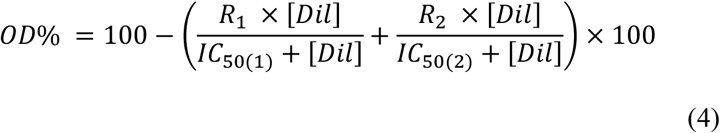

All analyses and graphs were performed using Sigmaplot 10.0.

## RESULTS

### Development of the Humoral Response and Inhibition of PSC Motility

The immunization protocol (Fig. 1) induced a progressive humoral response in llamas, as evidenced by motility and viability assays (Fig. 2). Sequential serum collections after each inoculation enabled longitudinal evaluation of antibody development and its functional consequences.

HIS produced a marked inhibition of PSC viability (Fig. 3) and a dose-dependent reduction in parasite movement (Fig. 4), with minimal effects at high dilutions and near-complete suppression at high serum concentrations. Exposure of PSCs to HIS resulted in clear morphological and functional alterations, accompanied by loss of viability (Fig. 3). Representative micrographs (Fig. 2, left panels) show intact PSCs under control conditions (Fig. 2, left, top) and structurally compromised organisms following exposure to immune serum (Fig. 2, left, bottom), as evidenced by methylene blue uptake.

**Fig. 3:**
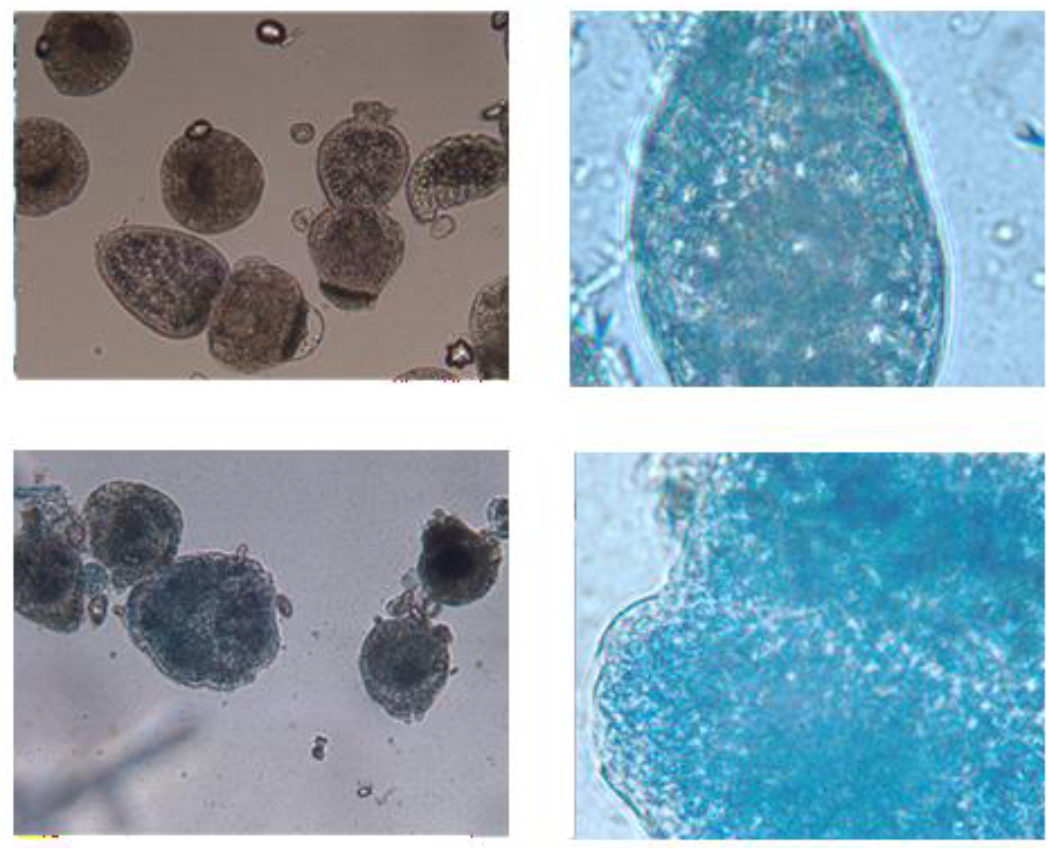
Effect of HIS and NS on Morphology and Viability of PSCs. HIS resulted in clear morphological and functional alterations (Bottom panels), accompanied by loss of viability as evidenced by methylene blue uptake, while NS showed no effect (Top panels).

**Fig. 4:**
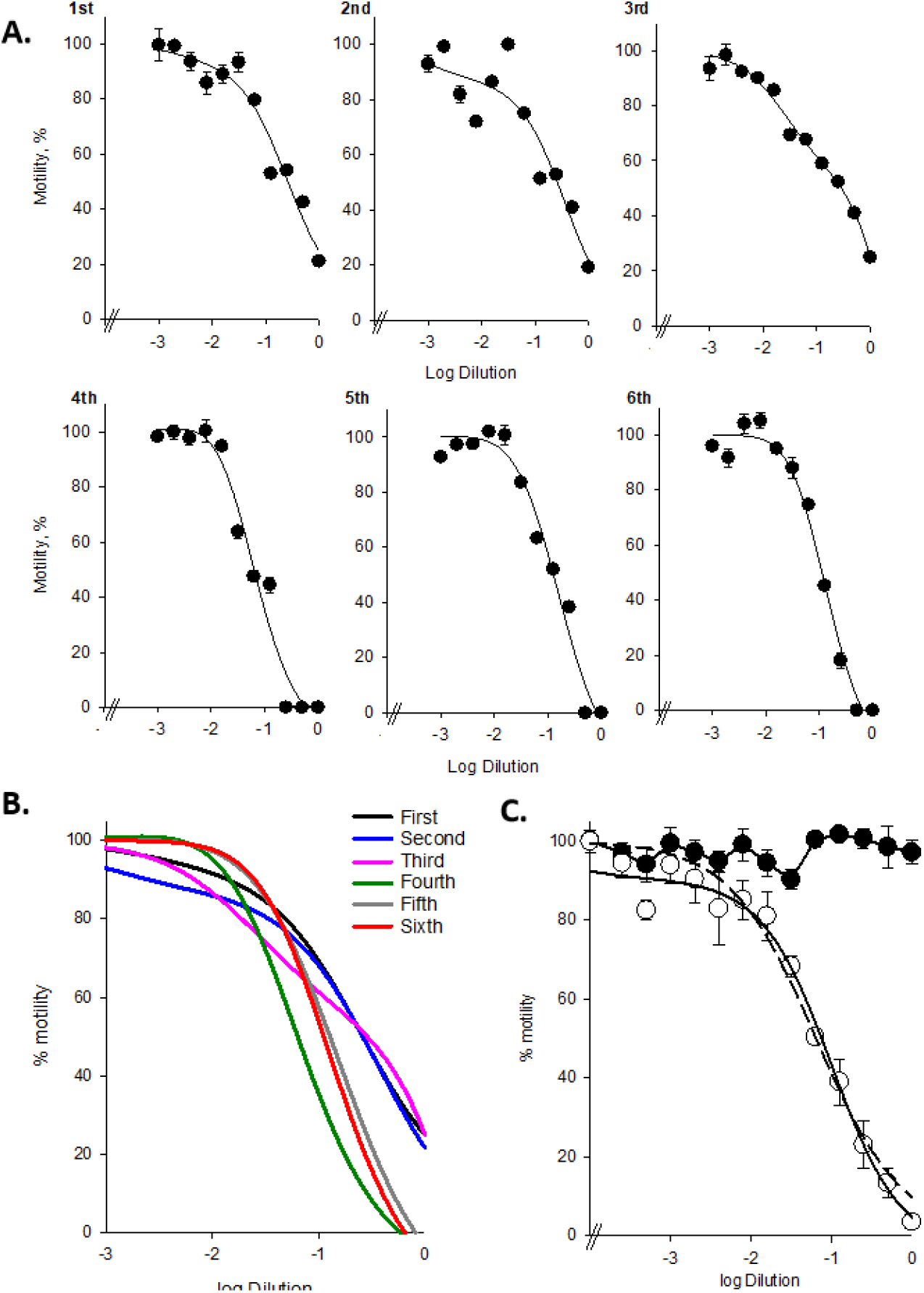
Dose-dependent inhibition of PSC motility by hyperimmune serum. **A**. Individual dose– response curves of PSC motility obtained with sera collected after successive immunizations (1st– 6th). Each panel shows nonlinear fits illustrating a progressive increase in inhibitory potency across immunizations. **B**. Superimposed curves highlighting the leftward shift in IC_50_ and increased slope associated with antibody maturation. **C**. Representative average response comparing HIS after 6^th^ inoculum (open circles), and naïve serum (filled circles). Dashed lines indicate Hill model fits, whereas solid lines represent two-site model fits, demonstrating improved agreement at intermediate dilutions.

Naïve serum-maintained motility close to the control value. These results indicate that antibody-mediated effects impair a key physiological function of the parasite even at the time of initial injection.

Analysis of data from sera after successive immunizations, fitted to the two-site model, revealed a progressive enhancement of inhibitory activity (Fig. 4). Early sera (1st–2nd inoculations) exhibited shallow curves and weaker inhibition. In contrast, later sera showed a leftward shift in IC_50_ (Table 1), steeper slopes (greater cooperativity), and stronger maximal inhibition, consistent with affinity maturation and/or expansion of antibody populations with improved functional activity.

**Table 1.** Chronology of inoculation response on PSC motility.

|  | Parameters |  |  |  |  |
| --- | --- | --- | --- | --- | --- |
| | $IC_{50(1)}$ | $IC_{50(2)}$ | $R_1$ | $R_2$ | $r^2$ |
| **1<sup>st</sup> inoculation** | 0.25 | 0.0024 | 0.8612 | 0.0636 | 0.9499 |
| **2<sup>nd</sup> inoculation** | 0.36 | 0.0007 | 0.8959 | 0.1265 | 0.8638 |
| **3<sup>rd</sup> inoculation** | 11.9 | 0.024 | 4.2245 | 0.4336 | 0.9883 |
| **4<sup>th</sup> inoculation** | 0.046 | 0.011 | 1.6191 | -0.5107 | 0.9697 |
| **5<sup>th</sup> inoculation** | 0.15 | 0.0096 | 1.3090 | -0.1049 | 0.9744 |
| **6<sup>th</sup> inoculation** | 0.061 | 0.0607 | 122.8 | -121.6 | 0.9866 |
| **Average** | 0.100 | 0.0000272 | 0.94 | 0.095 | 0.9900 |

### Evidence for Heterogeneous Inhibitory Response. The Two-Site Model

While the Hill model adequately described the overall sigmoidal behavior, systematic deviations were observed, particularly at intermediate dilutions (Fig. 5). To address this, the data were fitted with a two-site model for individual llama fits, and model comparison was performed assuming independent components with distinct apparent affinities. Titer–response curves for selected serum samples (e.g., IDs 713 and 720), showing fits obtained with both Hill (dashed lines) and two-site (solid lines) models. Parameter estimates (apparent affinities and slopes) and goodness-of-fit metrics (*r*^2^) are indicated. The two-site model consistently improves the fit, supporting the presence of multiple functional binding components.

**Fig. 5:**
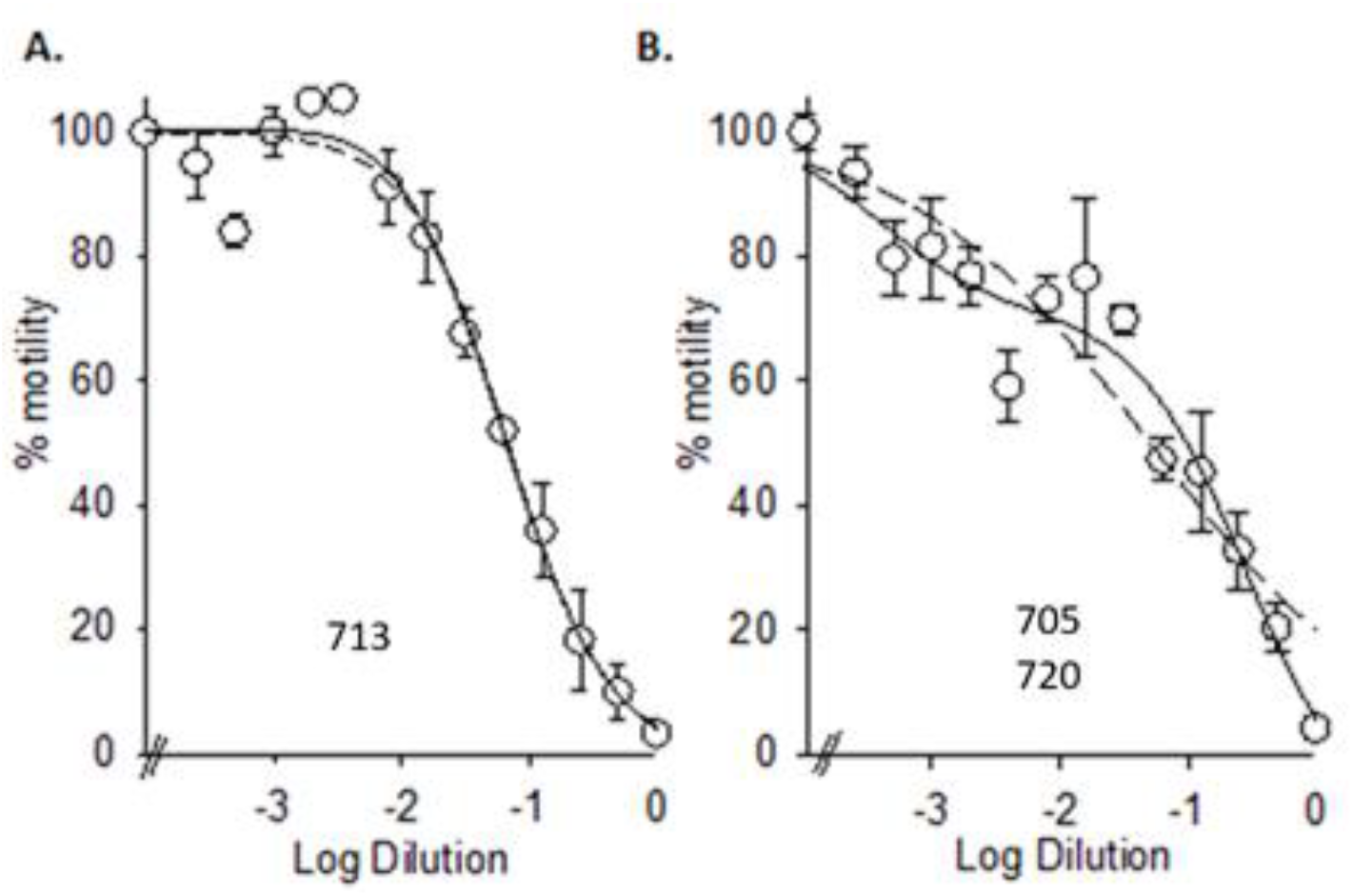
Model comparison of inhibitory responses. Dose–response curves for representative sera, e.g., llama IDs 713 (panel **A**), and IDs 705-720 (panel **B**). Data were fitted using both Hill (dashed line) and two-site (solid line) models. The two-site model provides improved fitting, particularly at intermediate dilutions, supporting the presence of heterogeneous antibody populations with distinct apparent affinities.

This approach provided improved fits across the full dilution range (Table 2). The analysis revealed a high-affinity component (low *IC*_50(1)_, dominant at low dilutions) and a low-affinity component (higher *IC*_50(2)_, contributing at intermediate dilutions), consistent with a heterogeneous polyclonal antibody response targeting multiple epitopes or functional sites on the parasite.

**Table 2:** The top and Bottom parameters represent the maximum and minimum values. Log IC_50_, IC50, Hill Slope, and error IC50 of llama serum 713, 705, and 720.

| Llama ID | $IC_{50(1)}$ | $IC_{50(2)}$ | $R_1$ | $R_2$ | $r^2$ |
| --- | --- | --- | --- | --- | --- |
| 705-720 | 0.2905 | 0.0004 | 0.8464 | 0.2869 | 0.9549 |
| 713 | 0.0587 | 0.0039 | 1.1216 | -0.1002 | 0.9785 |

### ELISA for Titrating Anti-*E. granulosus* Antibodies in Individual Llamas

The serum from immunized camelids was stored at -70°C until processing. To determine the presence of specific IgG antibodies against Eg, an imported anti-llama antibody was kindly provided by Dr. Esteban Pingitore (IMMCA, CONICET-UNT), and Maxisorp ELISA plates were donated by IVEMA.

### Estimation of the Dissociation Constant (*K*_*d*_)

In this study, ELISA-based titration curves were generated using sera from three llamas immunized with an *E. granulosus* antigen (Fig. 6). The objective was to estimate the relative affinity of the generated antibodies by calculating the apparent dissociation constant (*K*_*app*_). The *K*_*app*_ represents the concentration of free antibody at which 50% of the antigen is bound. Under equilibrium conditions and with a signal proportional to the Ag-antibody complex, *K*_*app*_ can be approximated by the *IC*_50_ value obtained from the fitting to a sigmoidal curve (Fig. 6).

**Fig. 6:**
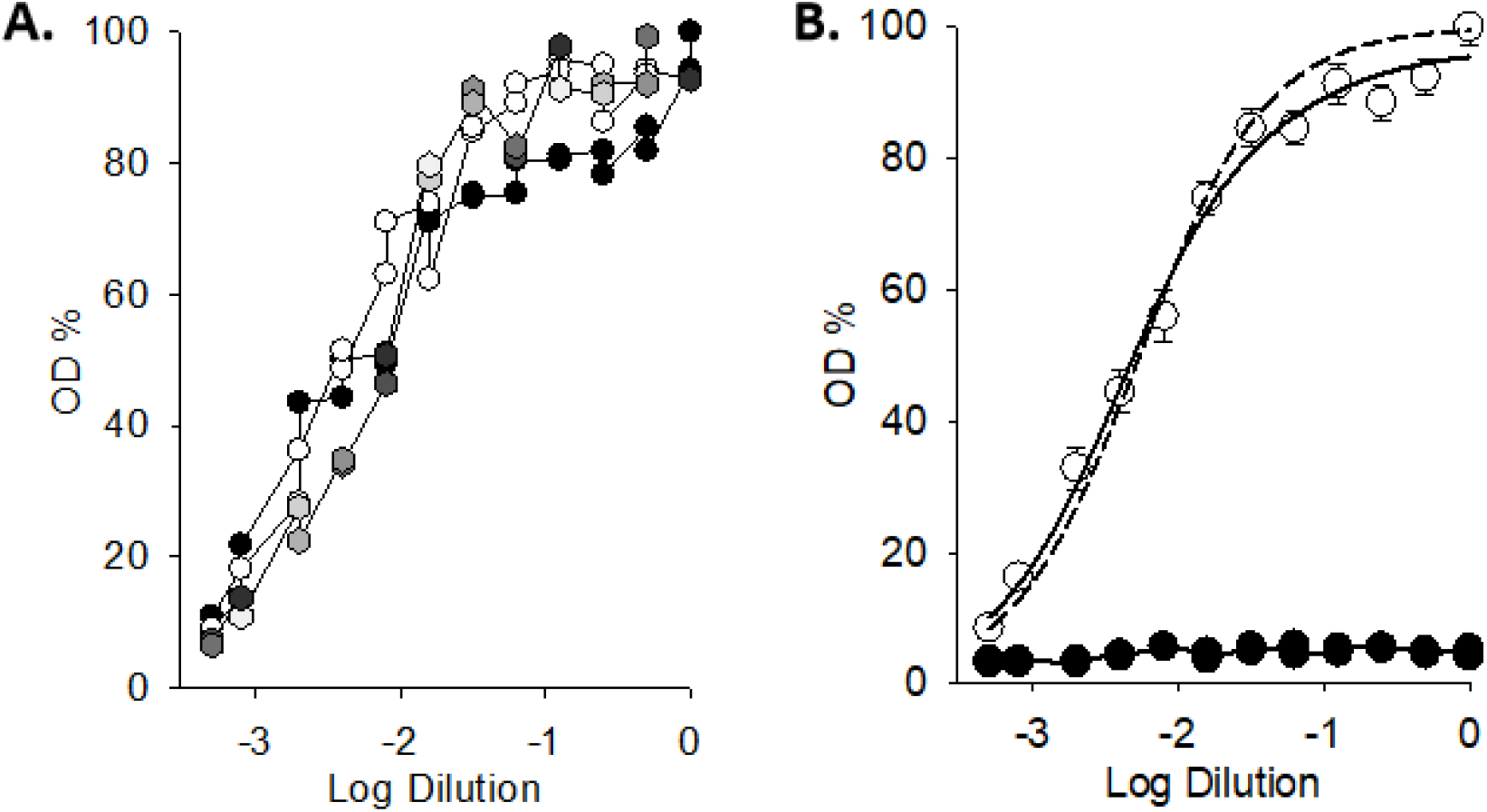
ELISA-based antibody titration and functional correlation. **Left:** Representative ELISA titration curves for sera from individual llamas. **Right:** Mean ELISA response for sera obtained after the sixth immunization. Data were fitted using both Hill (dashed line) and two-site (solid line) models.

**Fig. 7:**
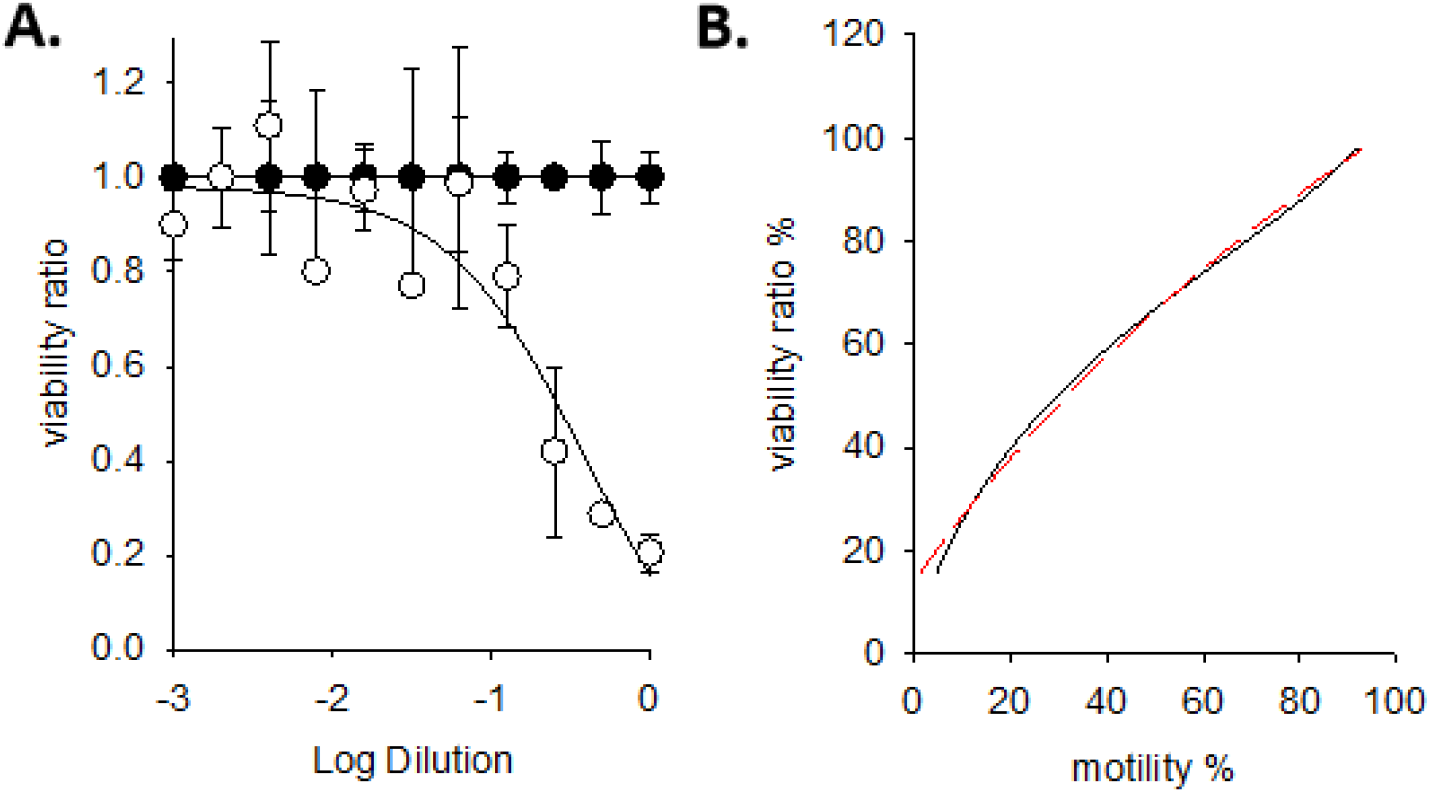
Comparative modeling and residual behavior. **A:** Viability data are shown for HIS and NS, in open and filled circles, respectively. Values are the Mean ± SD for at least three determinations. Fitting is shown with the two-site (solid line) model, which provides an improved description of the data, particularly in the intermediate dilution range. **Right:** Correlation between motility and viability (solid line) and the best fitting as shown with a red dashed line, demonstrating a strong correlation. The fitted line follows an exponential growth approach, however largely linear in this region, as f = y_0_ + a × (1 − e^(−b×x)^), y_0_ = 13.6, a = 151, and b = 0.0087, with an r^2^ of 0.9972, and SE of the estimate of 1.3166.

Assuming a total serum IgG concentration of 10 mg/ml (typical value in South American camelids), the following formula was applied:

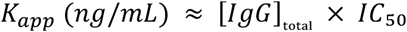

Additionally, *K*_*app*_ (Table 3) was calculated in molarity, considering a molecular weight of 150 kDa for IgG.

### Estimation of Specific Antibody Concentration

Based on the *IC*_50_ values obtained for each serum sample and assuming a total IgG concentration of 10 mg/ml (Life Diagnostics, Inc. 2018), the concentration of specific anti-*Echinococcus granulosus* antibodies (*AB*) was also estimated in each serum sample. This is based on the direct relationship:

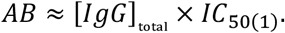

Since *IC*_50(1)_ represents the dilution at which 50% of antigenic sites are occupied, it can be considered a functional estimate of the level of specific antibodies in the original serum. The *IC*_50(1)_ represents the concentration of free antibody at which 50% of the antigen is bound. Mathematically, it is defined as:

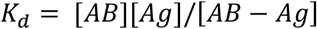

Under equilibrium conditions and with a signal proportional to the amount of *Ag-AB* complex, *K*_*d*_ can be experimentally approximated by the *IC*_50(1)_ value obtained from equation (4).

Assuming a total serum IgG concentration of 10 mg/mL (typical value in South American camelids), the *K*_*d*_ can be estimated as follows:

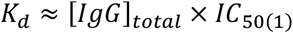

For example, for the mean ELISA response, whose *IC*_50_ was 0.1000:

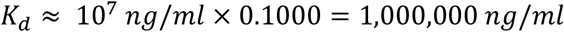

This value can be converted to molarity if an estimated molecular weight of 150 kDa for IgG is considered:

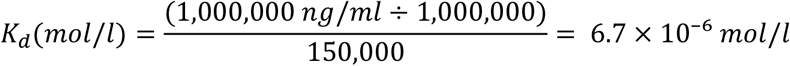

This procedure allows comparison of the relative affinity of antibodies between treatments. However, it should be noted that the results obtained represent an apparent constant (*K*_d,app_) when dealing with polyclonal sera with mixtures of affinities (Table 4).

For a precise determination of *K*_d_, techniques such as surface plasmon resonance, bio-layer interferometry, or competitive ELISA with purified antibodies would be required.

## Discussion

The present study demonstrates that hyperimmune sera generated in llamas against *E. granulosus* antigens exert a robust and reproducible inhibitory effect on protoscolex physiology, as evidenced by a parallel reduction in motility and viability. These effects are strongly concentration-dependent, antigen-specific, and progressively enhanced across successive immunizations, supporting the development of a functionally active humoral response with increasing potency.

A central finding of this work is the tight coupling between motility inhibition and loss of viability. Across the full range of serum dilutions, both parameters followed similar sigmoidal dose–response relationships and exhibited a strong positive correlation (r = 0.86). This indicates that motility impairment reflects a genuine loss of biological integrity rather than a reversible or sublethal effect. This observation extends classical viability criteria in protoscolex assays, where motility and dye exclusion have traditionally been considered complementary but independent indicators (Smyth & Davies, 1974; Smyth & Barrett, 1980). In contrast, our results demonstrate that, under antibody-mediated conditions, motility constitutes a reliable quantitative surrogate for parasite viability.

This concordance is consistent with previous reports in drug-based protoscolicidal assays, where motility loss precedes or accompanies irreversible structural damage (Elissondo 2006; Ceballos 2015). Our findings therefore support the use of motility as a sensitive functional readout of parasite integrity. While recent studies from our group have shown that pharmacological modulation of ion channels and cytoskeletal components can differentially affect motility and viability (Carabajal 2026 in print), the present results suggest that antibody-mediated inhibition leads to a more tightly coupled functional collapse of both processes.

From a quantitative perspective, the dose–response behavior of both motility and viability was initially well described by a Hill-type model, consistent with a cooperative inhibitory process characterized by a defined IC_50_ range. However, systematic deviations from this model, particularly at intermediate serum dilutions, revealed a more complex interaction landscape. This was especially evident in certain samples (e.g., llama 713 compared to 703 and 720), where the single-site model failed to adequately capture the full response profile.

To address this, the data were analyzed using a two-site model, which consistently improved the fit across datasets. This model revealed the presence of at least two functionally distinct inhibitory components with different apparent affinities. A high-affinity component appears to dominate at low dilutions, producing rapid functional inhibition, whereas a lower-affinity component contributes to the overall effect at intermediate concentrations.

Biologically, this multi-component behavior is consistent with the expected heterogeneity of polyclonal antibody responses and the structural complexity of the parasite tegument. It likely reflects the simultaneous targeting of multiple epitopes with distinct functional roles, including those involved in tegumental integrity and neuromuscular activity. The progressive emergence of these components across successive immunizations, as evidenced by leftward shifts in IC_50_ and increased curve steepness, is consistent with affinity maturation and expansion of antibody populations with enhanced functional relevance.

The functional impact of hyperimmune serum is further supported by morphological evidence of tegumental disruption and structural collapse of protoscoleces following exposure to high serum concentrations. Previous studies have demonstrated that *E. granulosus* PSCs can activate complement in vitro, leading to measurable changes in tegumental physiology (Ferreira 1992). Although electrophysiological measurements were not included in the present study, the parallel decline in motility and viability strongly suggests that immune recognition at the parasite surface translates into profound physiological disruption.

The relationship between ELISA titers and functional inhibition further reinforces the biological relevance of the humoral response. Sera exhibiting higher antigen-specific reactivity consistently showed greater inhibitory potency, indicating that ELISA-derived parameters can serve as predictors of biological activity. This relationship is particularly valuable for the selection and optimization of immunobiological reagents, as it directly links molecular recognition to functional outcomes.

Camelids produce both conventional IgG and heavy-chain–only antibodies, the latter giving rise to single-domain fragments (nanobodies) with unique biophysical properties, including high stability and the ability to access cryptic epitopes (Hamers-Casterman et al., 1993; Muyldermans, 2013). Although the present study does not distinguish between antibody isotypes, the strong inhibitory effects observed suggest that camelid antibodies are particularly well suited to targeting complex parasite surfaces such as the tegument of *E. granulosus*. This raises the possibility that the functional activity described here could be further refined by isolating specific nanobody populations with enhanced specificity and potency.

From a broader perspective, these findings are consistent with previous evidence indicating that antibody responses can interfere with parasite survival and development in cestode infections (Rogan & Richards, 1986; Heath et al., 1996). By demonstrating that hyperimmune llama serum can directly impair PSC motility and viability in vitro, this study provides functional support for developing passive immunization strategies or antibody-based interventions targeting the early stages of infection.

Despite these advances, several limitations should be considered. First, the experiments were conducted in vitro using a limited number of animals, and thus, the extent to which these findings translate to in vivo conditions remains to be established. Second, the contribution of complement and other serum components was not directly evaluated, and future studies should aim to dissect the relative roles of antibody-mediated and complement-dependent mechanisms. Third, although the two-site model provides an improved quantitative description, it remains a phenomenological representation, and further work will be required to identify the specific molecular targets underlying each component.

In conclusion, hyperimmune sera generated in llamas against PSCs of *Echinococcus granulosus* induce a strong, concentration-dependent impairment of parasite function, characterized by tightly coupled inhibition of motility and viability. The quantitative analysis reveals a multi-component inhibitory response consistent with polyclonal antibody heterogeneity and affinity maturation. These findings validate motility as a sensitive functional endpoint and highlight the potential of camelid-derived antibodies as versatile tools for developing novel immunotherapeutic strategies against cystic echinococcosis.

